# Scalable graph analysis tools for the connectomics community

**DOI:** 10.1101/2022.06.01.494307

**Authors:** Jordan K. Matelsky, Erik C. Johnson, Brock Wester, William Gray-Roncal

## Abstract

Neuroscientists now have the opportunity to analyze synaptic resolution connectomes that are larger than the memory on single consumer workstations. As dataset size and tissue diversity have grown, there is increasing interest in conducting comparative connectomics research, including rapidly querying and searching for recurring patterns of connectivity across brain regions and species. There is also a demand for algorithm reuse — applying methods developed for one dataset to another volume. A key technological hurdle is enabling researchers to efficiently and effectively query these diverse datasets, especially as the raw image volumes grow beyond terabyte sizes. Existing community tools can perform such queries and analysis on smaller scale datasets, which can fit locally in memory, but the path to scaling remains unclear. Existing solutions such as neuPrint or FlyBrainLab enable these queries for specific datasets, but there remains a need to generalize algorithms and standards across datasets. To overcome this challenge, we present a software framework for comparative connectomics and graph discovery to make connectomes easy to analyze, even when larger-than-RAM, and even when stored in disparate datastores. This software suite includes visualization tools, a web portal, a connectivity and annotation query engine, and the ability to interface with a variety of data sources and community tools from the neuroscience community. These tools include MossDB (an immutable datastore for metadata and rich annotations); Grand (for prototyping larger-than-RAM graphs); GrandIso-Cloud (for querying existing graphs that exceed the capabilities of a single workstation); and Motif Studio (for enabling the public to query across connectomes). These tools interface with existing frameworks such as neuPrint, graph databases such as Neo4j, and standard data analysis tools such as Pandas or NetworkX. Together, these tools enable tool and algorithm reuse, standardization, and neuroscience discovery.

## Introduction

Nanoscale connectomics researchers commonly need to analyze connectome networks that now surpass the computational power of single consumer workstations. As multimillion-edge connectomes become more commonplace (see Fig. 1), scalable and distributed computing approaches are needed to efficiently query and analyze these networks. The connectomics community has begun to develop tools to make this process easier and more efficient, either by subsampling networks to process subsets independently (1), or by adopting existing industry standards for graph storage and processing (2).

**Fig. 1.**
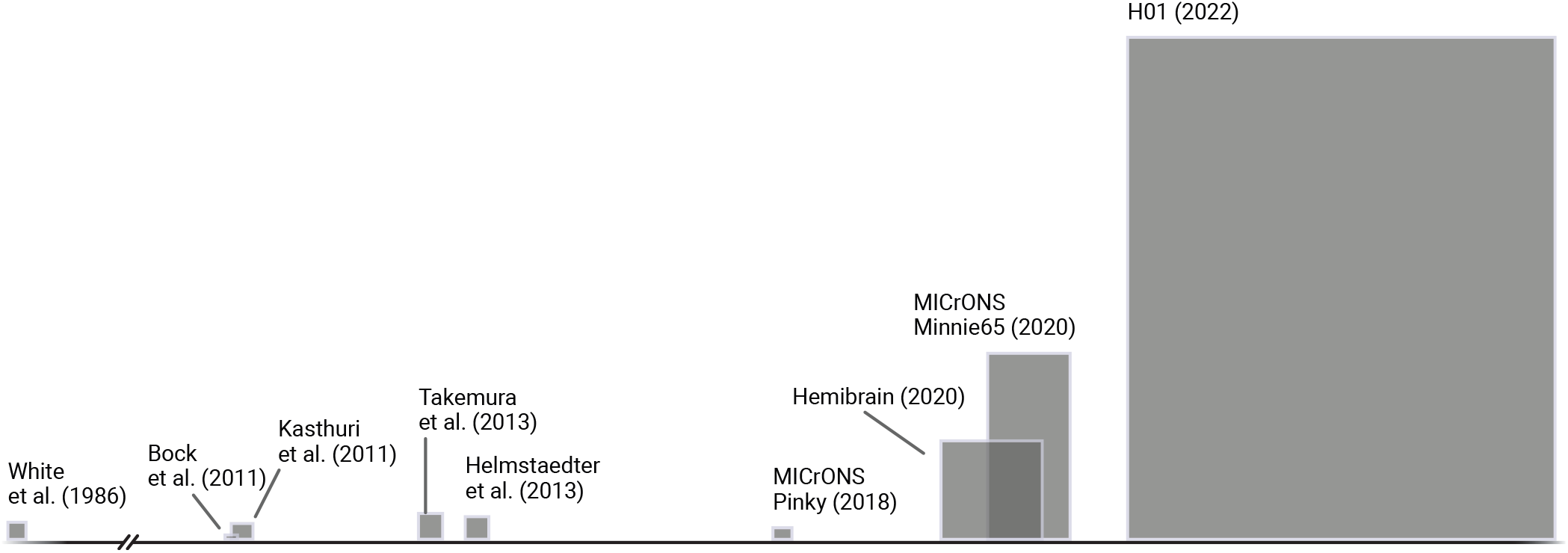
A non-exhaustive comparison of connectome network sizes. Graphs are arranged along a horizontal time scale. Widths are scaled to be proportional to the number of unique synaptic targets in the dataset, and heights are scaled to be proportional to the number of unique synaptic sources in the dataset. Both proofread as well as unproofread neurons (that may be errorful) are included in these numbers. Isolate vertices and edges are omitted. At this scale, the square representing the complete mouse connectome would be approximately the width of this page. Statistics used to generate this figure were computed with *Grand* and are available in **Table 4**.

While the total size of modern datasets continues to grow, the subset of immediately relevant data is often much smaller than the full dataset (1, 3). This is both a technical challenge as well as logistical challenge for many researchers, who must first query the full dataset to find the relevant subset, and then analyze the subgraph. Long-running, persistent tools like graph databases are a powerful option to enable high speed graph queries, but these databases may incur a significant maintenance cost even in the absence of complicated, demand-based scaling. Non-database approaches, like storing on-disk edgelist files, are also useful, but require more expertise to write distributed graph analysis workflows. Both approaches tend to require significant technical expertise, which is not always available to the broader neuroscience community. Ongoing neuroinformatics efforts in this space seek to create tools for analysis of these large connectomics datasets. Examples include efforts to create new relational data models and processing pipelines, and dataset specific query engines. An important gap that we address in this work is a suite of community tools to enable scalable, comparative analysis.

We share several approaches to query and analyze large-scale connectome networks while ensuring that operational costs and logistical burden remains low. We describe these tools, and share our experience and recommendations for the future of cloud-scale connectomics and network neuroscience.

## Contributions

In this work, we present a suite of tools for cloud-scale connectome analysis and graph queries. We assert that as the connectomics community grows and continues to ask increasingly complex questions of the nervous system, tools like these will enable high-throughput and high-impact neuroscientific discovery.

We present MossDB, Motif Studio, Grand, and GrandIso-Cloud, and explain how these tools fit into the greater connectomics analysis ecosystem, which can be found at github.com/aplbrain.

### MossDB

Data science workflows often use an immutable source-of-truth database to store products, as a way of tracking data provenance. We developed MossDB to serve as an associative datastore for relating graph data, imagery data, and other annotations. MossDB is a simple schemaless wrapper API around industry standards like MongoDB or DynamoDB (or even flat JSON files on disk) that provides (1) immutable references to data, without storing the data itself, (2) authentication, and (3) dataset discovery for secondary analysis.

MossDB was designed to be used alongside spatial datastores like BossDB (4) or CloudVolume (5), and graph databases like Neo4j (6) or Grand (discussed in further depth below). Because the datasets in question can span hundreds to thousands of terabytes, it is not realistic to store copies of even small subsets of the data. Instead, MossDB uses URI-based linking in order to reliably point to dependable and immutable datasets on these remote services.

When an administrator sets up a new MossDB instance, they can allow a set of URI prefixes to which MossDB users can link. For example, the URI prefix-set of the MossDB instance used for the analyses in this paper is shown in **Table 1**. Validators — written in Python — can be attached to each of these types to ensure that data exist at the specified endpoint when a new MossDB document is created.

**Table 1.**
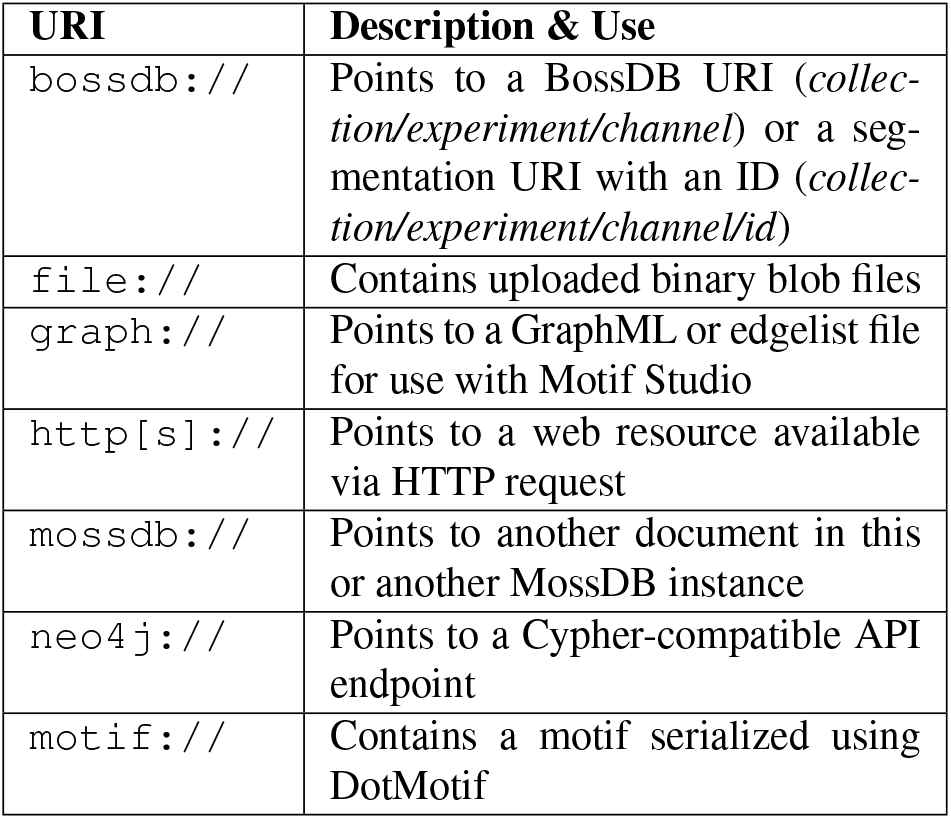
A list of URI prefixes supported by the MossDB instance used to conduct the analyses in this paper. Additional user-specified prefixes may be added through a minimal alteration to the motif-studio runtime configuration.

**Table 2.** A summary of selected backends available in the Grand Python package. Backends handle data storage and graph operations, and are independent of the API with which a user interacts with a dataset.

| Backend | Description & Notes |
| --- | --- |
| DataFrameBackend | Stored in Pandas dataframes |
| DynamoDBBackend | Edge/node tables in DynamoDB |
| GremlinBackend | Gremlin graph databases |
| IGraphBackend | An IGraph graph, in memory |
| Neo4jBackend | Neo4j graph databases |
| NetworkitBackend | A Networkit graph, in memory |
| NetworkXBackend | A NetworkX graph, in memory |
| SQLBackend | Edge/node SQL-queryable tables |

**Table 3.** A summary of dialects available in the Grand Python package. Dialects can be used to interact with graph data through a familiar API, regardless of the underlying data storage backend.

| Dialect | Description & Notes |
| --- | --- |
| CypherDialect | Neo4j-like Cypher API |
| IGraphDialect | IGraph-like interface |
| NetworkXDialect | NetworkX-like interface |
| NetworkitDialect | Networkit-like interface |

**Table 4.** Connectome sizes from **Figure 1**. Note that as connectome network sizes trend upward, network density decreases considerably. Numbers here are reported for proofread as well as unproofread data. Synaptic and neural isolates are omitted. Datasets are ordered by the number of unique source neuron IDs. Note that the number of unique targets, and total graph density, do not correlate monotonically with the unique number of source neurons. Directed edge density is computed as 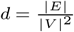, where |*E*| is the number of unique directed edges in the dataset and |*V* | is the number of unique vertices in the dataset.

|  | Unique sources | Unique targets | Density | Reference |
| --- | --- | --- | --- | --- |
| Bock et al. | 11 | 185 | 0.005657 | Bock et al. (8) |
| MICrONS Pinky v185 | 104 | 334 | 0.015608 | (9–11) |
| White et al. | 276 | 296 | 0.039246 | White et al. (12) |
| Kasthuri et al. | 380 | 857 | 0.001607 | Kasthuri et al. (13) |
| Helmstaedter et al. | 1015 | 1075 | 0.458212 | Helmstaedter et al. (14) |
| Takemura et al. | 1471 | 1155 | 0.010612 | Takemura et al. (15) |
| Hemibrain v1.2 | 21692 | 21738 | 0.007513 | Xu et al. (16) |
| MICrONS Minnie65 | 106126 | 13009 | 0.000250 | (9–11, 17) |
| H01 | 1268187 | 794401 | 0.000001 | Shapson-Coe et al. (18) |

Because authentication and data access needs vary by research group, MossDB has a flexible authentication system that can be easily swapped for alternatives based upon the needs of the userbase. The simplest supported authentication model is secret-based authentication (where a user uses a passphrase when writing documents, and must reuse that passphrase in order to update or delete those documents), but this can be swapped for more complex authentication systems, such as OAuth, as use-cases require.

Finally, it is important that such a tool enables dataset discovery for secondary analysis. MossDB URIs are indexed by — and can be searched by — unique prefixes, which make it easy to identify annotations for a given dataset (i.e., all annotation documents attached to the *bossdb://witvliet2020/Dataset_1/em* hierarchy) without incurring major string-matching computational expense. To aid in data exploration, we provide a simple HTML web interface for browsing MossDB projects (**Figure 2**).

**Fig. 2.**
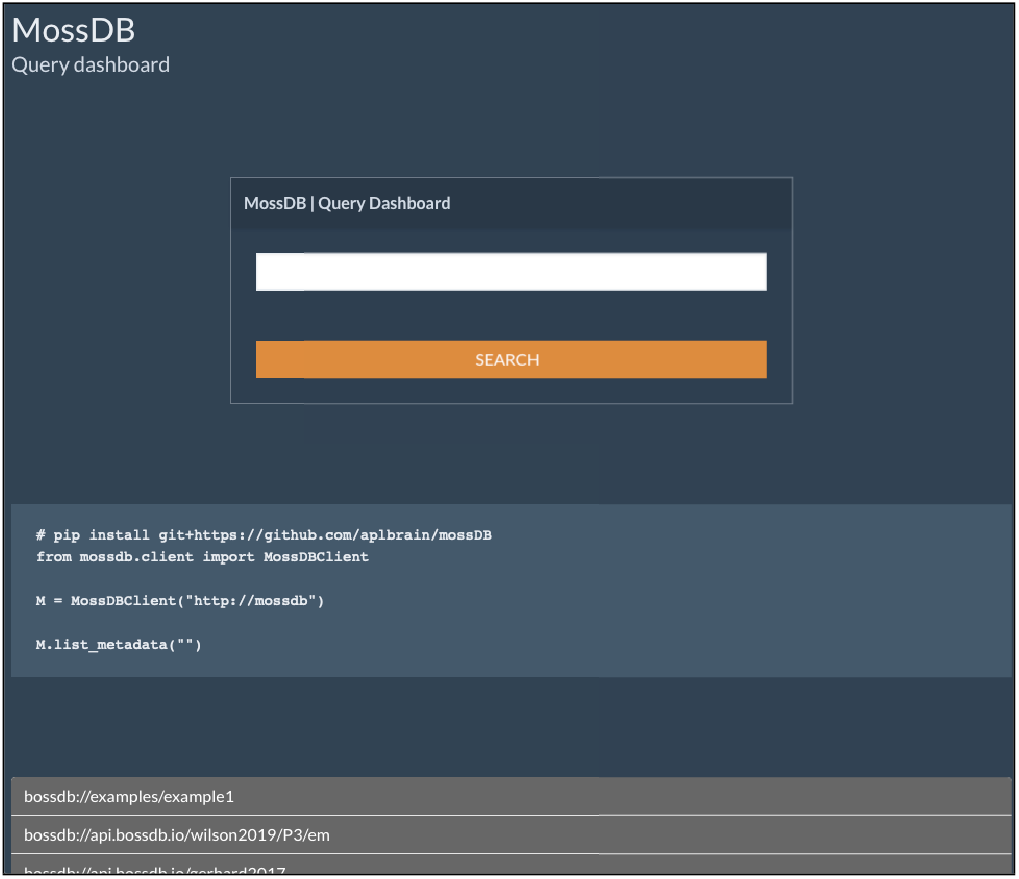
A simple MossDB web interface, illustrating the functionality to explore information across multiple datasets. Code to reproduce the UI search query is generated automatically, in order to encourage reproducible analysis.

The source code and documentation for MossDB are available online at https://github.com/aplbrain/mossDB.

### Motif Studio

While DotMotif (Matelsky et al. (7)) dramatically lowers the barrier to entry for new graph motif searches, its use still requires a familiarity with the Python interpreter and common graph file formats. (7) In order to accelerate hypothesis-testing and scientific discovery, the community needs accessible, zero-configuration tools.

Therefore, we developed Motif Studio, a browser-based query-design environment that provides real-time visual feedback to users as they curate and develop graph queries (**Figure 3**). Motif Studio is written in Python Flask (for graph management, search, and HTTP API) with a React TypeScript front-end user interface, and can run in *server* mode (on a persistent, devoted compute node) or in *serverless* mode, in which case Motif Studio may be deployed to ephemeral infrastructure like AWS Lambda. In *serverless* mode, Motif Studio only incurs cost overhead when in active use.

**Fig. 3.**
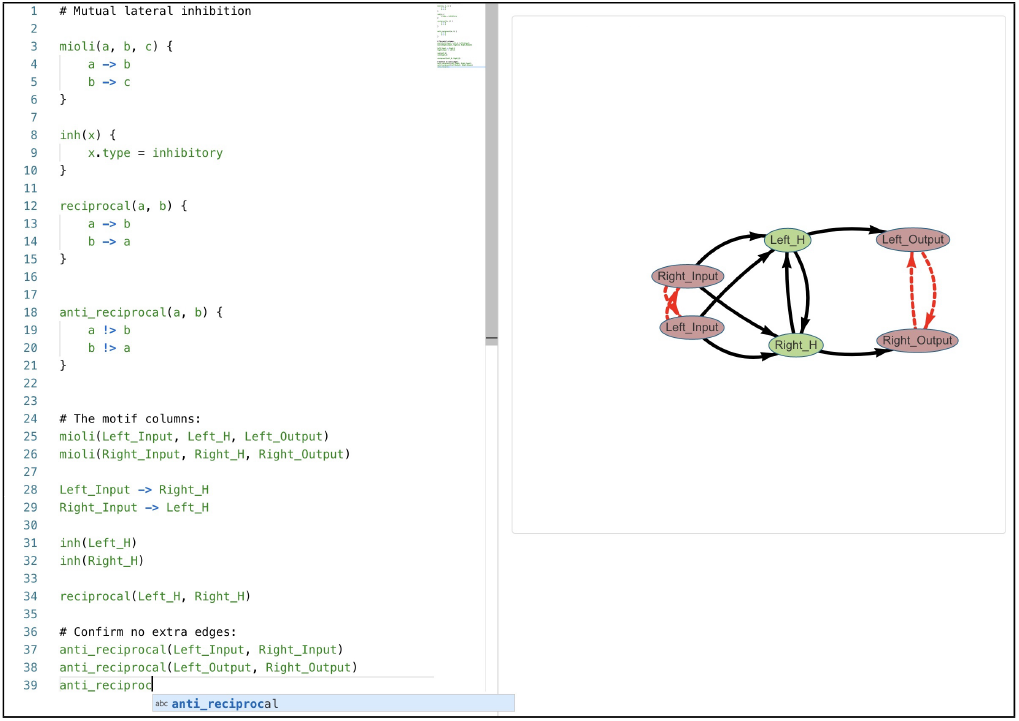
MotifStudio. A browser-based user interface for searching for motifs in connectomes. The tool is currently in *build* mode, and has side-by-side code-editor and motif-visualization panes. Query errors are detected in real time. The visualization pane and webpage URL are updated as the user types. The state of the application is encoded in the page URL, which may be shared with collaborators.

The architecture of the Motif Studio Python server is composed of a HTTP API server that points to a series of *Host-Provider* objects, which must satisfy a simple interface to enable graph searches. Motif Studio can serve as a query terminal — performing graph searches on its own captive compute — or as a query pass-through, leveraging Grand (described below) to relay queries to cloud datastores or other databases, such as neuPrint (2). Administrators of Motif Studio may also optionally permit end users to upload their own networks in common formats such as GraphML or edge-list CSVs.

#### Editing graph queries

Motif Studio has two primary windows — a query design window, and a query running window. In the editing view (**Figure 3**), the user can edit a motif description on the left using the DotMotif language (7).

On the right, the motif is visualized and updated in realtime. Nodes are colored by a unique hash of their attributes, which enables a query designer to see which nodes have like or disparate attribute constraints. Edges are colored black when they must exist in query results, and red when the user has specified that the edge must *not* exist in the host graph.

The code editor is equipped with syntax highlighting, based upon a DotMotif *tmLanguage* syntax file derived from the Backus–Naur grammar. This syntax highlighter, and a Microsoft Visual Studio Code extension for editing DotMotif .*motif* files, is available at https://github.com/aplbrain/dotmotif-vscode/. The editor also supports text autocomplete, multiple cursors, and token matching.

#### Performing graph queries

Graph queries can be performed on local compute (best for small graphs or user-uploaded graphs) or relayed to higher-powered compute resources, such as Neo4j servers or neuPrint HTTP APIs (2, 6). Individual Motif Studio servers can also provide combinations of these options by supplying multiple *HostProvider* objects at runtime.

When running a graph query, the user may choose whether Motif Studio will return automorphisms and whether edge direction will be considered when searching. The former option is most useful when motifs or graph queries have a large number of symmetries; excluding automorphisms is an easy way to reduce the runtime of such a query. The edge direction option is useful when a general anatomy is hypothesized but synaptic direction is not yet known. This mode is also useful for testing motif prevalence in graphs that do not have explicit directions attached to each edge (e.g., MRI brain networks).

Results of the executed query are shown in the bottom panel, where each vertex of the motif is assigned to a column, and each valid mapping (i.e., monomorphism) of the motif is returned as a unique row in the results pane table. Execution-time errors, if any, are shown to the user in the results pane or build pane, depending on the provenance of the error.

Motif Studio can tightly integrate with the MossDB metadata store, which enables a Motif Studio user to leverage companion datastores such as annotation servers or imagery datasets. For example, the official BossDB Motif Studio deployment uses deep links in the results pane to bring a user to a Neuroglancer visualization of the participants in a motif, if segmentation is available in BossDB. Each update to the motif query, or changes to the selected host graph, also updates the web UI URL in realtime. These Motif Studio URLs can be shared with peers or collaborators to work cooperatively on a research question.

A publicly accessible version of Motif Studio is running at https://motifstudio.bossdb.org.

### Grand

Algorithm reuse is a critical component of the scientific process. Grand is a Python library that maps scalable graph operations to familiar, non-scalable APIs. In other words, a developer can reuse algorithms written in NetworkX or IGraph to operate on graphs much larger than would fit in RAM on a conventional consumer workstation. This capability to transparently scale existing work is particularly impactful when graph storage in RAM is a primary constraint on runtime, which is common in graph traversals and queries. Grand comprises three primary components — *Dialects*, an operation abstraction layer, and *Backends* (**Figure 5**). The end-user interacts with Grand through one or more *Dialects*, which are sets of API calls designed to emulate (or inherit from) well-known, industry-standard libraries. For example, the *NetworkXDialect* is a set of functions designed to look like the commonly-used NetworkX Python library (19). The user may then choose a *Backend* tool that will actually perform read- or write-operations on the network itself. This pattern is inspired by tools such as Dask or Modin, which maintain compatibility with familiar library APIs while introducing additional infrastructure that improves runtime performance (20–22).

**Fig. 4.**
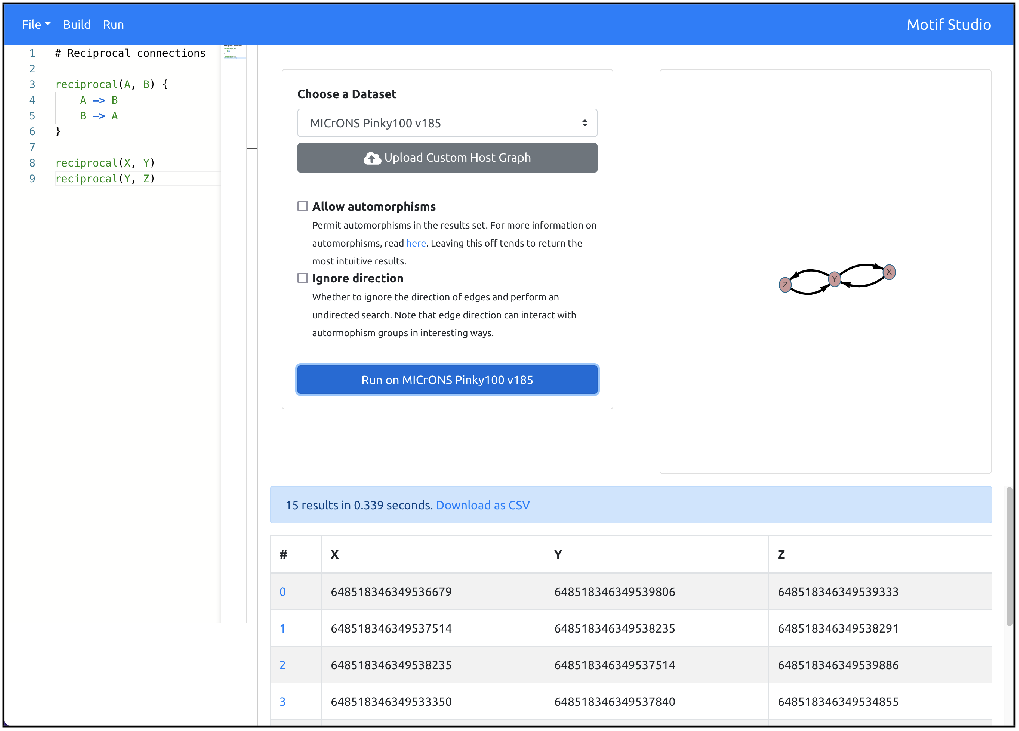
MotifStudio. A browser-based user interface for searching for motifs in connectomes. The tool is shown in *run* mode. The code editor and visualization panes from Figure 3 are still visible. A host-graph selection screen is also visible in the middle, and results are rendered in the bottom panel.

**Fig. 5.**
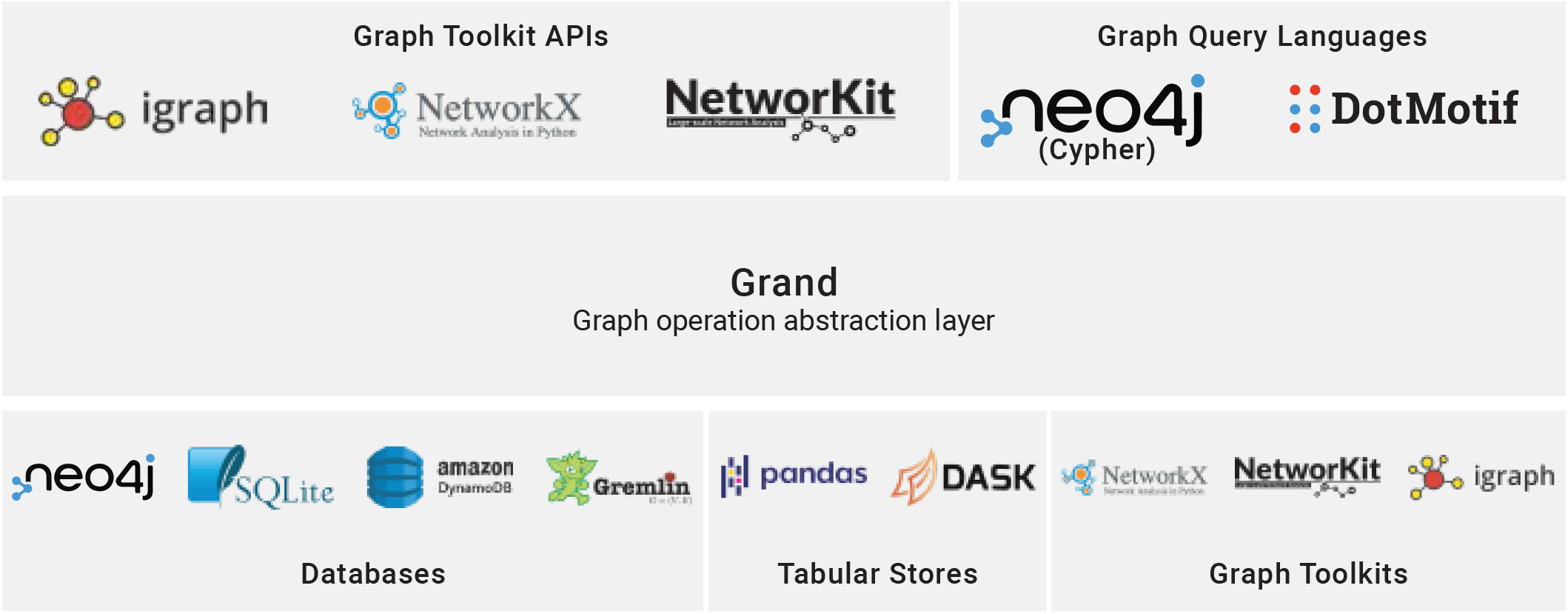
The Grand graph library architecture. Grand supports multiple *dialects*, or input syntaxes (top row). Commands from these dialects are converted into an abstract internal representation for generalized graph manipulation and queries. The end user may then select a *backend* in which to store the network and run the operations themselves. Grand supports common graph toolkit API dialects — such as NetworkX — to enable researchers to rapidly prototype and then scale algorithms to larger compute resources. Grand also supports common graph query languages like Cypher (Neo4j), and the domain-specific motif-query language, DotMotif. These dialects enable researchers to smoothly interoperate between cloud-scale big-data datasets and local, small scale networks. Grand supports delegating workloads to various backends, including commonly used databases such as SQLite; tabular datastores, such as the popular Pandas Python library; or graph toolkits themselves. This interoperability enables write-once/run-anywhere algorithm design for the network science research community.

The ability to combine dialects with backends means that a user can prototype an algorithm for a small, in-memory NetworkX graph, and then run it at scale on a performant SQL database by using the *NetworkXDialect* and the *SQLBackend*. As a concrete example of this portability, we adapted the GrandIso algorithm first described in Matelsky et al. (7) to run on cloud-scale, multi-million-edge networks by using Grand. The resulting algorithm, *GrandIso-Cloud*, is described below.

### GrandIso-Cloud

Subgraph monomorphism, the underlying algorithm behind motif search, is memory-hungry, and most state of the art implementations may require the allocation of many terabytes of RAM even for relatively small host graphs (thousands to tens-of-thousands of edges). The GrandIso algorithm isolates this memory cost in a single, one-dimensional queue data structure. In the original GrandIso tool (written for the NetworkX Python library (19)), the queue resides in memory (7). This makes it extremely fast, but it also means that the total size of the motif search task is limited by the RAM of the machine. This is a particular issue for modern large-scale graph analysis questions, where the host graphs may exceed hundreds of millions of edges. Such a graph would exceed the memory- and time-budgets of most institutions. Thus, there are two main limitations to consider: (1) The size of the raw graph data may exceed what can be stored on a single machine; (2) The RAM requirements of the queue may exceed the RAM of the machine.

For this reason, the GrandIso algorithm is an ideal candidate for scaling using the Grand tools described above. The GrandIso-Cloud implementation of the GrandIso algorithm outsources queue management to a dropout-resilient queue system like AWS SQS. (23) This has three main advantages: First, it eliminates local RAM limits as a bottleneck; the queue can scale exponentially on the remote cloud host without impacting local performance. Second, it enables multiple parallel workers — on the same machine or on multiple machines — to work cooperatively on the same graph, without requiring complicated parallelism or memory management mutexes. And finally, this persistent, out-of-memory queue adds a layer of resilience to drop-out or node death, which enables the user to recruit much larger clusters of unsupervised worker nodes.

Besides the obvious cloud-scale financial considerations, GrandIso-Cloud is to the best of our knowledge scalable to extremely large host network sizes. As we continue to refine and optimize this codebase, we expect that cloud-native network analysis will increasingly take place in near-realtime, collocated with the original spatial (imagery and segmentation) storage.

The source code for the GrandIso-Cloud tool is available at https://github.com/aplbrain/grandiso-cloud.

## Conclusions

Algorithm and dataset reuse are two foundational pillars of the scientific process, and the rich opportunity for secondary analysis depends on interoperable software and tools. Because of the fast-paced forward progress of the connectomics field in the past few years, tool interoperability amongst the community continues to grow in importance. It is critical to build connections and tools that encourage interfaces and innovations between community solutions. We hope that users interested in comparative connectomics and motif search can leverage these tools to accelerate discovery and hypothesis testing.

## Supplemental Materials

Tools are available online at github.com/aplbrain.

## ACKNOWLEDGEMENTS

The authors thank the creators of the connectome datasets discussed here. This work was supported in part by National Institutes of Health (NIH) grants R24MH114799, R24MH114785, and R01MH12668.

